# Winter Survival of *Culex pipiens f. pipiens A*dults in Central Greece

**DOI:** 10.1101/2024.11.06.622242

**Authors:** Charalampos Ioannou, Stavroula Beleri, Persa Tserkezou, Antonios Michaelakis, Eleni Patsoula, Christos Hadjichristodoulou, Nikos T. Papadopoulos

**Affiliations:** Department of Agriculture, Crop Production and Rural Environment, University of Thessaly, Magnisias, Greece; Department of Public Health Policy, School of Public Health, University of WestAttica, Athens, Greece; Laboratory of Hygiene and Epidemiology School of Medicine, University of Thessaly; Scientific Directorate of Entomology and Agricultural Zoology, Benaki Phytopathological Institute, Attica, Greece

**Keywords:** winter survival, longevity, mosquitoes, population dynamics, vector borne diseases

## Abstract

Winter survival consists a major component of insect vectors life history in temperate environments that is directly related with early and later population growth next season with major consequences in the epidemiology of vectored diseases. The common European mosquito *Culex pipiens* is a major vector of the West Nile Virus (WNV) in Europe, including Greece. West Nile Virus outbreaks are frequently reported in Greece over the last 2 decades and Thessaly, Central Greece, is included in the affected areas. Here we report on overwintering trials conducted in three regions of Thessaly to investigate the overwintering dynamics of the subspecies of the *Cx. pipiens* complex, *Cx. pipiens f. pipiens*. Two overwintering experiments regarding adults of *Cx. pipiens f. pipiens* carried out in two coastal areas of Thessaly (Nea Anchialos and Volos) and an inland area (Kalamaki). Results demonstrated the successful overwintering of *Cx. pipiens f. pipiens* females, as well as the failure of males to survive in all three regions considered. Successful overwintering females were capable of initiating egg laying following a blood meal in spring onsetting the first summer generation. Nonetheless, mortality patterns differ between the coastal and the inland area as well as among different cohorts of adults.

## Introduction

In regions with a temperate or cold climate, like Greece, mosquitoes have developed a variety of overwintering mechanisms that, depending on the species, may include the egg, larval, adult or more than one developmental stage [1–4]. Several factors, most importantly low temperatures and precipitation are those that determine the duration of the overwintering period and may vary for a given species depending on latitude [5–8]. Usually, winter temperatures in cooler temperate areas do not allow breeding and population growth of mosquitoes. This fact, combined with the high mortality rates observed during the cold months may result in a dramatic decline in populations that onset the first spring generation. From an epidemiological point of view, the proportion of a vector mosquito population that will successfully overwinter is of particular importance, as the initiation and development of the spring generation will depend on it [5]. Moreover, the overwintering of mosquitoes’ vectors is important not only for their population biology but also for the evolvement of associated diseases [3,9]. For example, the persistence of certain pathogens in overwintering mosquitoes may contribute in maintaining the transmission cycle each year, rendering the disease endemic [10,11].

West Nile virus (WNV, family: Flaviviridae) is currently the most important mosquito-borne pathogen spreading in Europe [12,13]. Data on overwintering of WNV in mosquitoes are crucial for understanding WNV circulation [14]. In temperate regions, most species of mosquitoes are subject to facultative diapause initiated by a decrease in day length and temperature, which results in the interruption of transmission cycles of mosquito-borne pathogens during winter [2,9]. The mosquito *Culex pipiens* (Diptera: Culicidae) is of growing concern, as it is considered the main vector of WNV in Europe [6,11,15] including Greece [16–18]. This species includes two distinct forms, known as “*pipiens*” (Linnaeus, 1758) and “*molestus*” (Forskål, 1775), which can form hybrids [7,19,20]. The two forms are morphologically identical but display important differences in their behaviour and physiology. One of the major distinctions between the biotypes is their overwintering biology [19,21–23].

In cold, temperate regions, the *’’pipiens’’* form overwinters as adult inseminated females entering into facultative reproductive diapause [2,6,24,25]. The factors responsible for the induction of female diapause are the low autumn temperatures combined with reduced photophase affecting the last larval stages (3rd and 4th) and the pupa [26,27]. In contrast, males do not enter diapause and do not survive the winter [23,27].

Dormant females are characterized by the absence of host search for blood meals and feed on “sugary” plant secretions, building up rich fat body reserves just before inhabit overwintering sites [27]. Feeding females with a 10% sugar solution for 7 to 10 days is sufficient to build up fat reserves [25]. Sites that remain frost-free during the winter such as caves, barns, underground storage facilities, channels and cracks in the ground are selected by adult mosquitoes as hibernating shelters. Although dormant individuals of *Cx. pipiens f*. *pipiens* do not exhibit host-seeking behavior, some females may be motivated and receive a blood meal when in close proximity to a host for a period of time [28], using the blood to build body fat rather than for ovarian development [29,30]. These blood meals are not useful to diapausing females either for body fat production or for ovarian development as they were found to be significantly inferior in quantity compared to non-diapause counterparts [25]. The ambient conditions (e.g. temperature, relative humidity) of the overwintering habitats may vary a great deal and determine female survival rates. For example, in cool and humid habitats with low fluctuation, females may remain at the same spot for weeks reserving precious stored energetic metabolites. In contrast, in more exposed to external conditions shelters, during the warmer hours of the day females may forage for sugar food or seek more appropriate overwintering places and activity that can risk survival and reduce longevity [1,2,6,9,19]. As soon as temperature increases in the spring, females abandon overwintering sites and forage for blood meals in appropriate hosts that assures egg maturation and oviposition.

In contrast to the above, *Cx. pipiens f. molestus* appears to be a taxon adapted to warmer climates and individuals of this form remain active during winter and can reproduce as long as temperatures allow (≥10 °C) in both surface and groundwater, mainly in groundwater [20,31,32]. At present*, Cx. pipiens* s.s. biotypes are regarded as distinct monophyletic evolutionary units [33–35]. As the biotype *“pipiens”* is usually found in aboveground habitats, while the biotype *“molestus”* is exclusively found in urban, below ground habitats, populations of both biotypes were considered genetically isolated in the northern regions [23 24,34,36–38]. However, despite their ecological and behavioural differences, *“pipiens”* and *“molestus”* occur sympatrically above ground in many European regions, and may interbreed to produce ‘hybrids’ where their distributions overlap [32,34,37,39].

We investigated the overwintering capacity of *Cx. pipiens f. pipiens* in three selected regions of Thessaly and in various habitats. We therefore aimed to quantify survival throughout the autumn and winter months, to define when they terminated diapause and start blood feeding in the spring and also to determine the impact of overwintering on vector competence of emerging *Cx. pipiens* mosquitoes for WNV.

## Materials and Methods

### Study areas and mosquito colonies

The overwintering experiments considering *Cx. pipiens f. pipiens* individuals were carried out in the area of Thessaly, central Greece, where three locations were chosen: (a) the village of Kalamaki on the mainland, adjacent to Lake Karla, (b) the town of Nea Anchialos, a coastal region and (c) Volos, the coastal port city of Thessaly (Fig. 1). Experiment 1 carried out in Nea Anchialos and Kalamaki in 2012-2013, while Experiment 2 took place in Volos, aiming to study in addition the possible effect of organic matter (as a food source) accumulating in the waters of the shelters during overwintering. In particular, overwintering adults were provided with either plain water (Treatment A) or plain water + organic water (a vial containing liquids/juices collected from a composting bin) (Treatment B).

**Fig 1.**
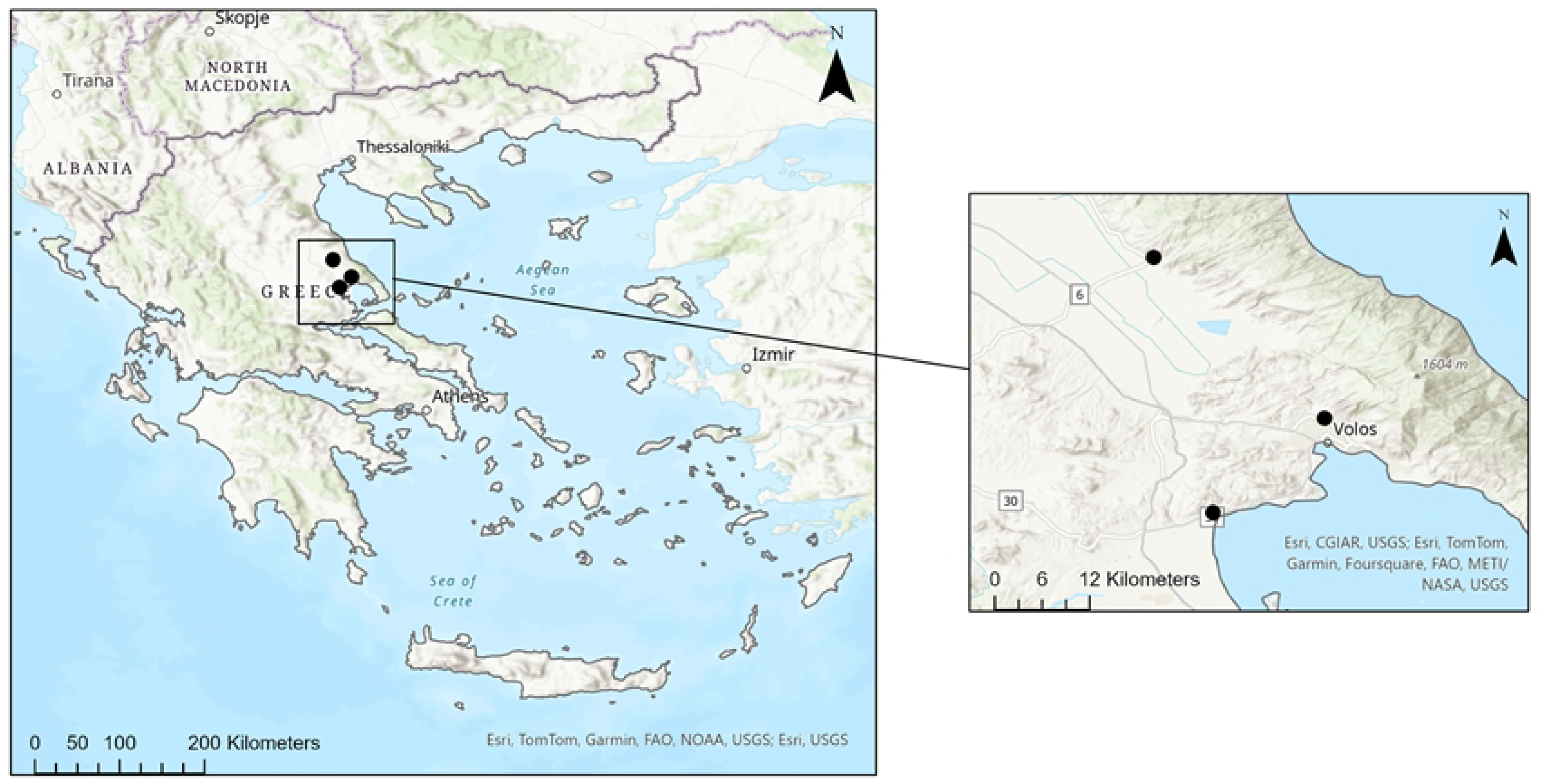
Map showing the experimental locations (Kalamaki, Nea Anchialos, Volos), where the experimental study took place.

The adult mosquitoes used in the overwintering experiments came from egg rafts that had been laid by laboratory-reared adults of *Cx. pipiens f. pipiens*. Colonization took place within the insectary of the laboratory of Entomology and Agricultural Zoology at the University of Thessaly, Greece.

The experimental procedure performed in this study included the following steps: hatching of *Cx. pipiens* eggs, development of immature stages to the adult stage, overwintering of adults in cages, which underwent three treatments, provision of blood meal to adult females and oviposition.

### Experimental procedures

The rafts of eggs (∼70) that were collected in the laboratory were transferred to a plastic container with 12 L of water and artificial food (Purina Adult cat food, Friskies) in a protected from rain outdoor sites of the Department of Agriculture, Crop Protection and Rural Environment at University of Thessaly, Volos, Greece.

During the development of the immature stages, both water and food were renewed at regular intervals, ensuring that the conditions were suitable until pupation. At the completion of development, 100 pupae, of both sexes, were transferred to plastic containers with 250 ml of water and placed in 20×20×20cm, Plexiglass cages until the emergence of adults.

Adults were transferred in a heated storage room (15 ± 2 °C) with natural lighting conditions and offered a 10% sugar solution during the first 10-12 days of life in order to build up the necessary adipose tissue reserves for overwintering and mating. After this period, the sugar solutions in the cages were replaced with plain water and then the cages were transferred to the overwintering sites. Five cages (500 adults in total) were transported to Kalamaki and Nea Anchialos on 30/12/2012 and 4/1/2013 respectively, and placed in moist and dark storage areas protected from rain and wind. Ten cages (1000 adults in total) that were placed to Volos on 1/12/2013 were randomly assigned to one of the following twotreatments: (a) five cages were provided with plain water (egg hatching 1/10/2013), (b) five cages with plainwater and plain water + organic water (egg hatching 1/10/2013). Treatment C was the same as Treatment A, entered later on the experimental procedure (egg hatching 15/10/2013) as a control.

In both experiments, at the end of the winter period (first two weeks of March), a 10% sugar solution was once again placed in the cages and then they transferred back to the semi-outdoor area of the Agricultural School at the University of Thessaly. This procedure took place on 7/3/2013 in Kalamaki, on 14/3/2013 in Nea Anchialos, and on 19/3/2014 in Volos.

The surviving (overwintering) females from Kalamaki and Nea Anchialos, were pooled in two cages and offered a two-hour blood meal via a special device for this purpose, on 11/4/2013. From 20/4/20213 to 22/4/2013 plastic containers with 250 ml of water were placed in the two cages (one from each overwintering location) for the females to lay their eggs. The overwintering females in Volos were provided with a blood meal on 23/4/2014 while the oviposition took place on 28/4/20214.

Survival of adults in the different treatments was recorded at regular intervals and dead individuals were removed from the cages. The deposited egg rafts were examined under the stereoscope to determine hatch.

### Meteorological data

The temperatures that prevailed throughout the experimental study, from the development of the immature stages to the oviposition of the females that survived in the three locations, were recorded with the help of special electronic devices (HOBO, Onset, USA), and are given in the following Figs 2-4. Figures include also information regarding the exposure period of adults in the overwintering sites as well as feeding and oviposition opportunities.

**Fig 2.**
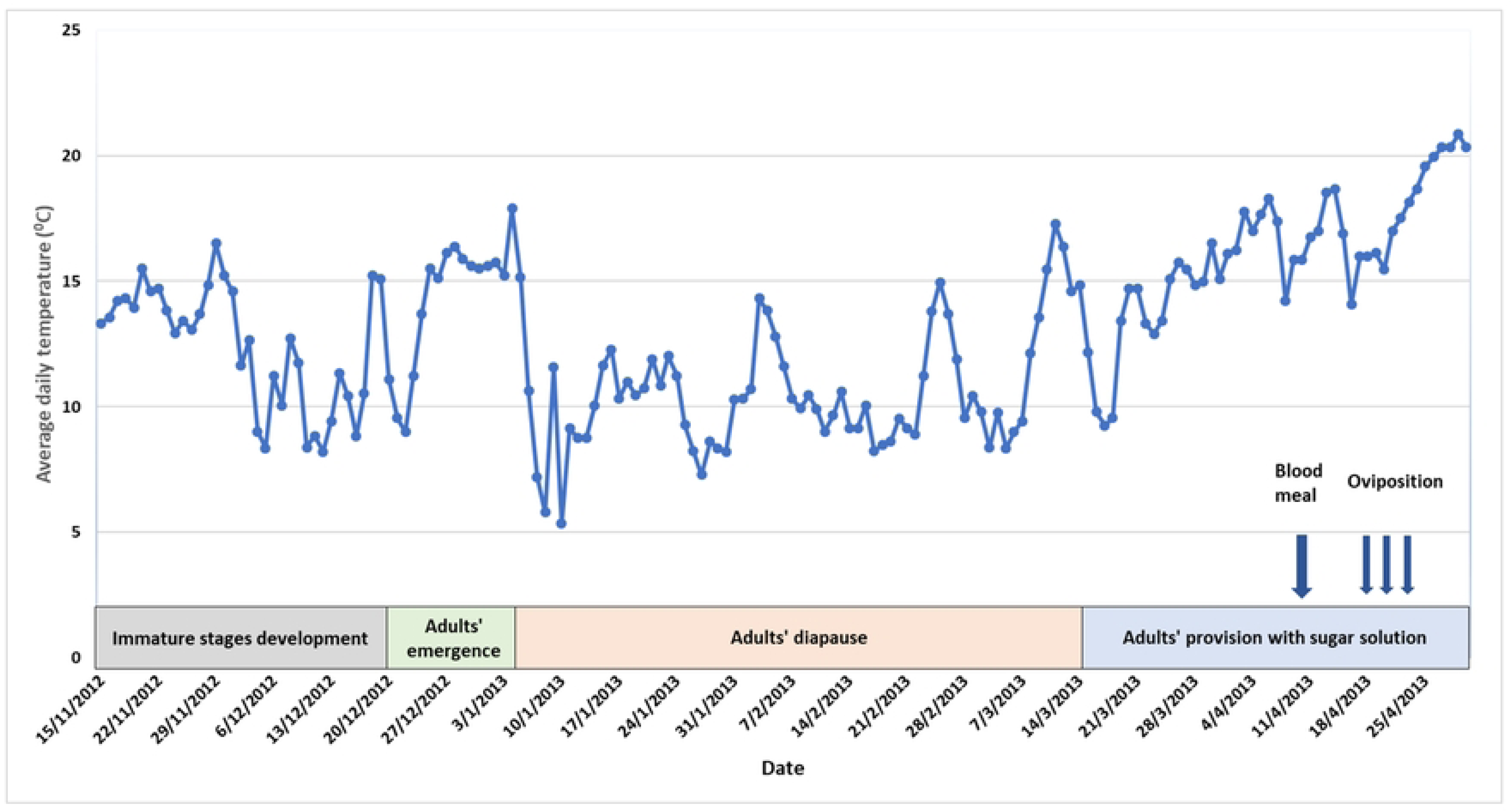
Average daily temperatures that prevailed throughout the experimental study of winter survival of *Cu/ex pipiens f pipiens* in Nea Anchialos (a coastal area) Thessaly, the period 2012-2013.

**Fig 3.**
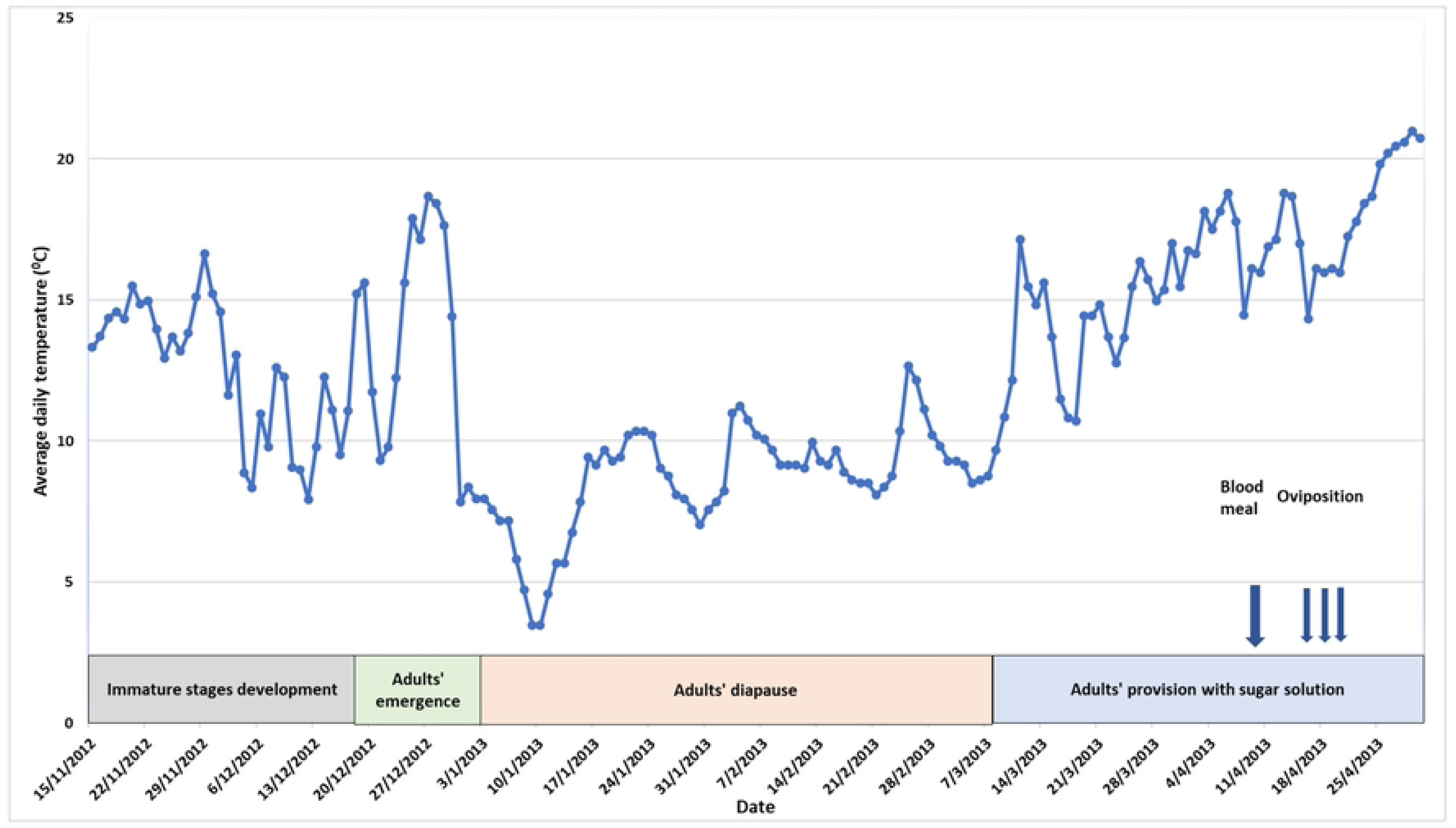
Average daily temperatures that prevailed throughout the experimental study of winter survival of *Cu/ex pip/ens f pip/ens* in Kalamaki (a mainland village), Thessaly, the period 2012-2013.

**Fig 4.**
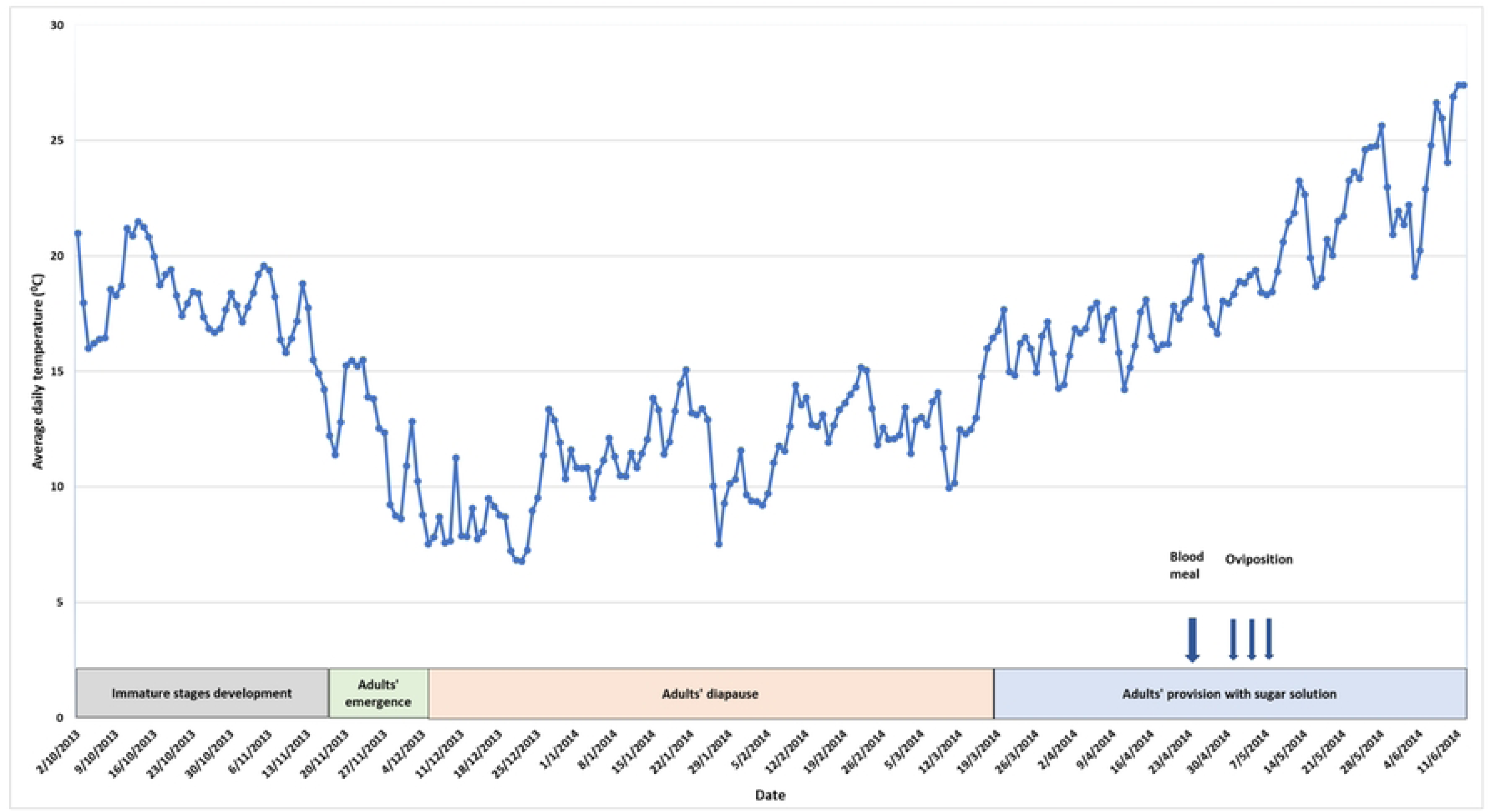
Average daily temperatures that prevailed throughout the experimental study of the winter survival of *Cu/ex pip/ensf. pip/ens* in Volos, Thessaly, 2013-2014.

### Statistical analysis

The Kaplan-Meier curves and the log-rank test was used to compare the survival times between males and females. Cox regression was employed to examine whether the sex of adults and the overwintering site were significant predictors of adult mortality rates. R version 4.3.2 (The R Foundation for Statistical Computing, Vienna, Austria) was used for data analysis. P values less than 0.05 were considered statistically significant.

## Results

### Winter survival of *Culex pipiens* f. *pipiens* in Kalamaki and Nea Anchialos

The average daily temperatures that prevailed in Nea Anchialos and Kalamaki during the experiment, from the development of the immature stages to the oviposition of the females that survived, are shown in Figs 2 and 3 respectively. In both areas temperatures were rather high until end of December and the cold period (winter) started in the beginning of January. Temperature increased in the beginning of March marking the end of winter period. Although comparable, winter temperatures were lower in the continental area of Kalamaki compared to Nea Anchialos. A total of 450 adults (90 per cage; 173 females, i.e. 38.95% of exposed individuals) were transported and exposed to winter conditions of Kalamaki, as determined at the end of the experimental procedure considering also accidental loses. The corresponding parameters for Nea Anchialos were 456 adults (91.2 per cages; 294 females, i.e. 64.65% of the exposed individuals). In both sites males did not manage to survive until the end of the cold season (early to mid-March) (S1 Fig). Higher survival rates for males were recorded in Kalamaki compared to Nea Anchialos. In Nea Anchialos, the mortality of females during the winter was progressive (S1 Fig). On the contrary, the survival of females in Kalamaki was particularly high until 26/2, followed by a significant decline after that date (S1 Fig). The addition of sugar solution with the rise in temperatures resulted in the stabilization of female mortality in both cases. At the start of oviposition (20/4) the average percentage of females that finally survived in Kalamaki and Nea Anchialos reached 23.85% (44 individuals in total) and 25.26% (76 individuals in total) respectively. Of these females, 52.27 and 47.37% oviposited respectively, while 95.65% and 91.67% of the laid egg rafts hatched.

Mortality rates of males and females at Kalamaki and Nea Anchialos were compared with the Kaplan Meir curves and the Cox proportion hazard model. Overall, and in both locations, male longevity was shorter than that of females with males failing to overwinter (Fig 5; *p* < 0.01). The hazard ratio of males compared to females was 12.9 (95%CI: 9.69, 17.2) in Kalamaki and 78.9 (95%CI: 47.4, 131.0) in Nea Anchialos.

**Fig 5.**
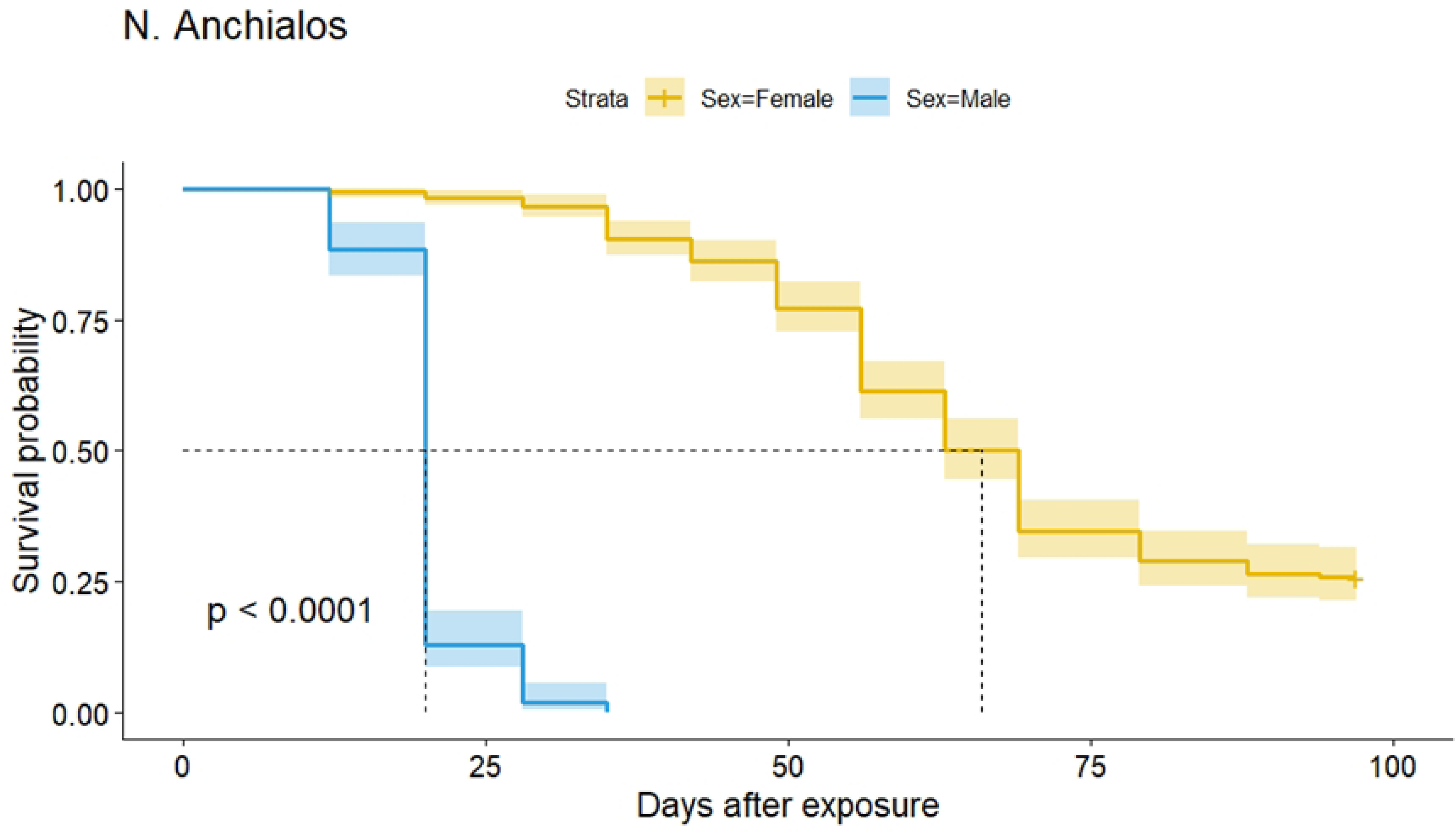
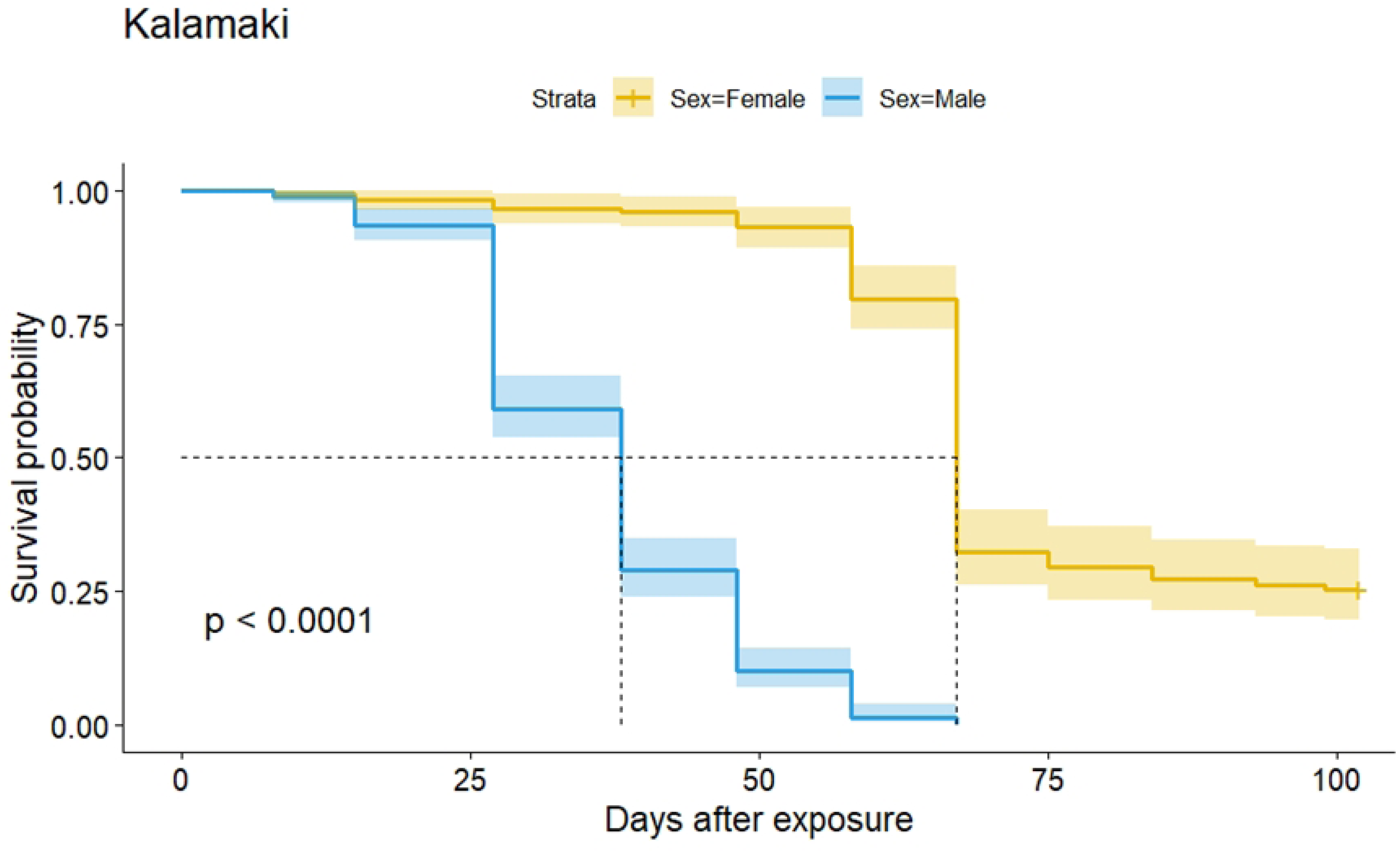
Kaplan Meier curves (with 95%CI) including log-rank test (shown p-value) for the survival of males and females in Kalamaki (A) and N. Anchialos (B). Dashed lines represent the median survival for males (20 days) and females (66 days).

Cox regression analysis considering location of exposure, sex and their interaction as predictors revealed that males have a significant higher hazard rate compared to females (*p* < 0.001), adjusted for location, and mortality rates were higher in Kalamaki compared to Nea Anchialos (*p* < 0.001) (Fig 6, S2 Fig). Overall hazard rates considering both males and females were similar between the two locations (*p* = 0.425). The significant interaction between sex and location is associated with higher mortality rates for males in Nea Anchialos compared to Kalamaki. Comparing survival patterns of females in the two location no significant differences were found (Fig 6; Log-rank test, *p* = 0.16).

**Fig 6.**
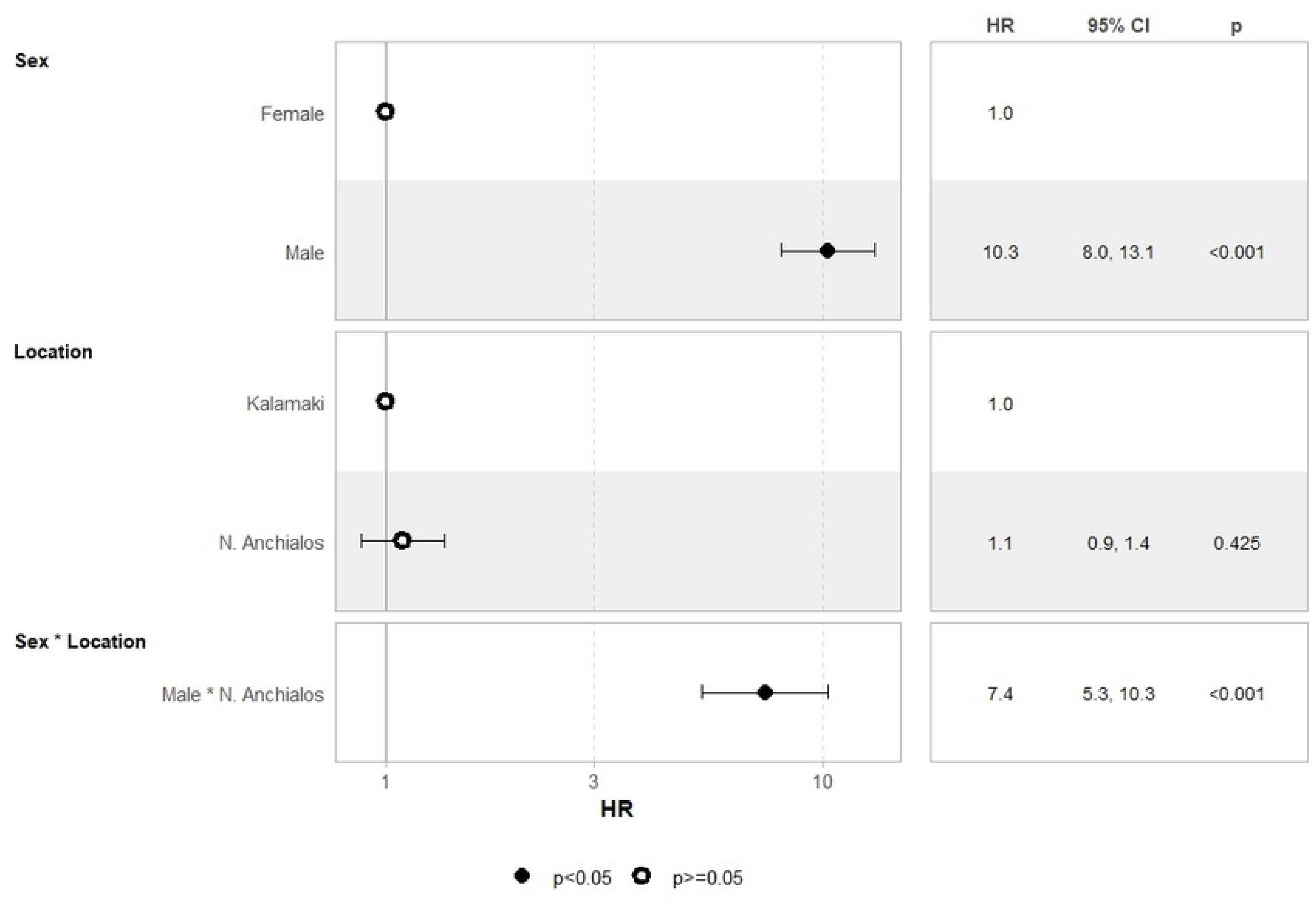
Forest plot depicting the hazard ratio for males andfemales exposed to Kalamaki and Nea Anchialos.

### Winter survival of *Culex pipiens f. pipiens* in Volos

The average daily temperatures that prevailed in Volos, during the overwintering experiment are shown in Fig 4. The temperature drop in the beginning of December marks the onset of winter while the increase in middle March the end. The total number of adults exposed to winter conditions was 482 (average per cage 96.4; 371 females, i.e. 76.9%), 475 (average per cage 95; 349 females, i.e. 73.5%) and 509 (average per cage 101.8; 388 females, i.e. 76.2%) for the treatments A, B and C respectively. In all three treatments male survival rates were much lower than that of females and no male survived after the end of December to reach the end of the winter period. Details of female and male survival patterns in each cage are given in S3 Fig. Among cage variation was low in treatment A and B and quite higher in treatment C. Overall, at the end of the winter period 9.42, 15.7 and 12.44% of the females managed to survive in treatments A, B and C respectively. Among cage variation in male survival was minimal within the same treatment and overall survival patterns among treatments negligible as well (S4Fig). Comparing the survival rates of the two sexes of *Cx. pipiens*, the ability of females (S5a Fig) to overwinter in all three treatments and the inability of males (S5b Fig) to survive was evident. In all three treatments provision of sugar solution on 16/3/2014 reduced mortality rates. At the start of oviposition (28/4) in treatment A, treatment B and treatment C, the number of females alive was 35, 53, and 48, of which 13, 25, and 10 oviposited, with the oviposition rate reaching 37.14, 47.17, and 20.83% respectively.

Kaplan Meir analysis followed by the long-rank test revealed the higher mortality of males compared to that of females within each treatment (Fig 7, *p* < 0.01). Cox regression analysis including treatment, sex and their interaction as predictors confirmed the overall higher hazard rates for males compared to females (*p* < 0.001) and differences among the three treatments (Fig 8). The hazard ratio of treatment B and C compared to baseline A was slightly but significantly lower, and higher respectively (Fig 8; *p* < 0.05). The interaction between sex and treatment was not significant. Kaplan Meir analysis comparing survival patterns of the three female groups followed by the log-rank test revealed significant differences among the three treatments (*p* < 0.01, Fig 7). Nevertheless, the proportion of surviving females at the end of the exposure period was similar among the three treatments (chi-square test, *p* > 0.05).

**Fig7.**
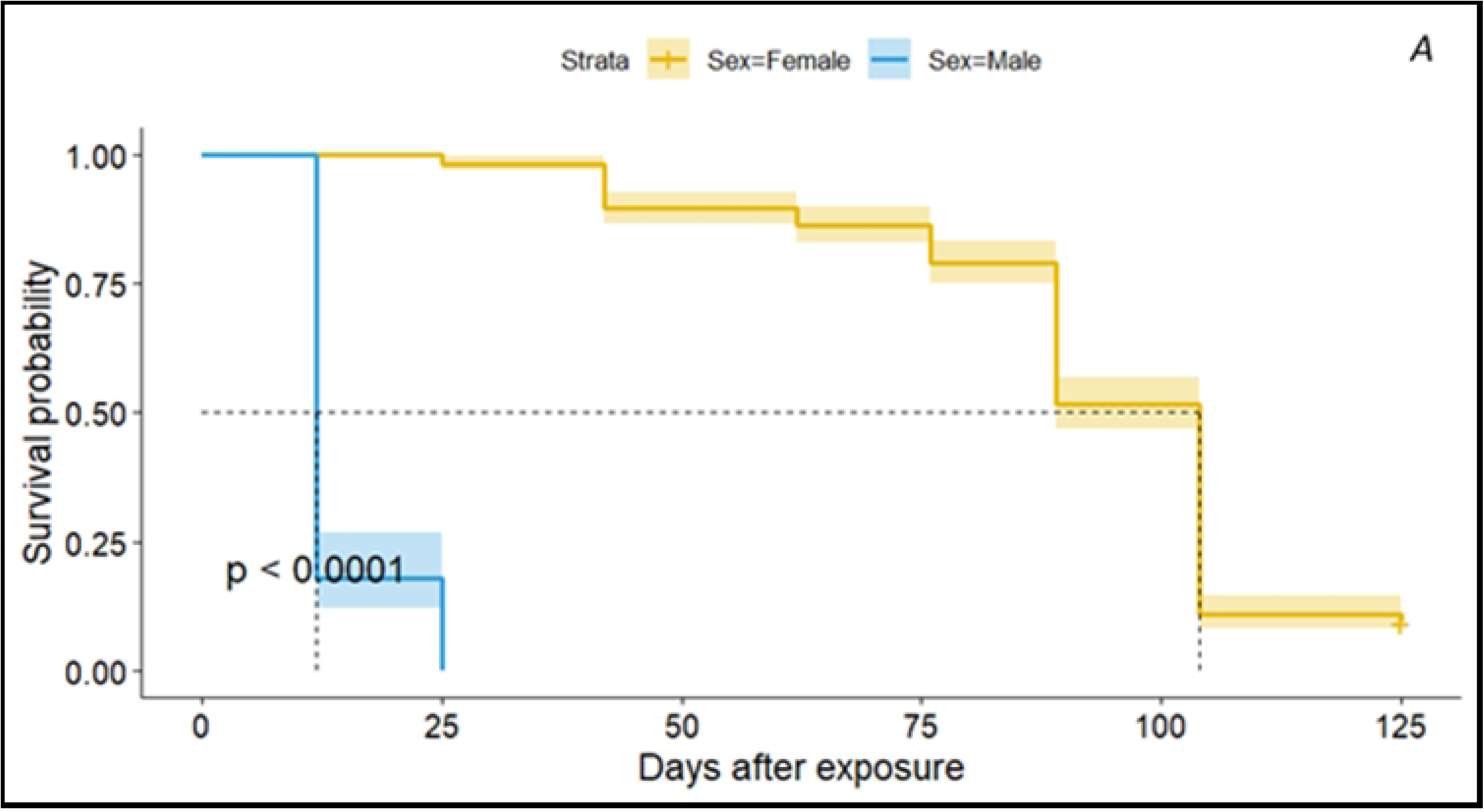
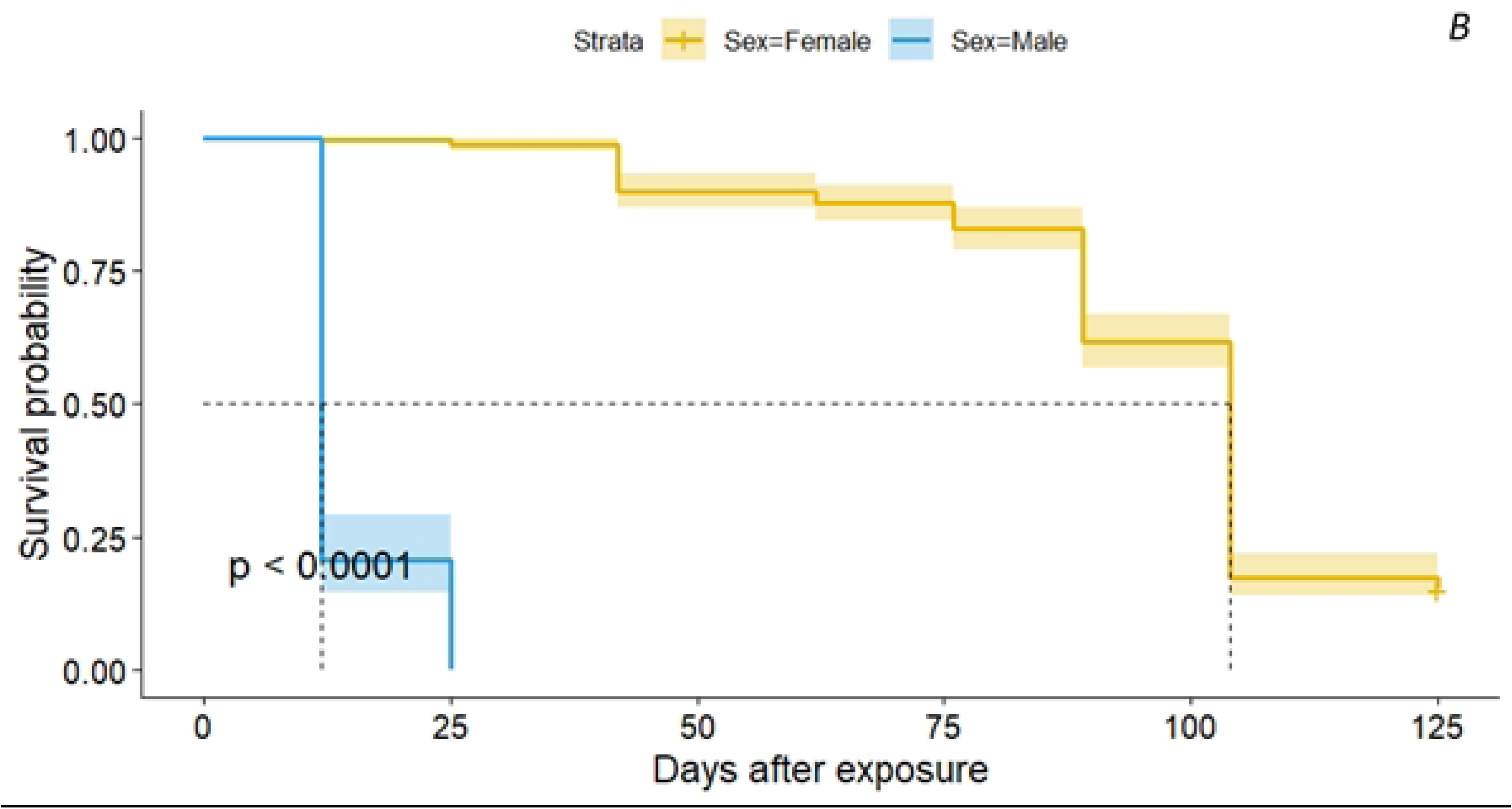
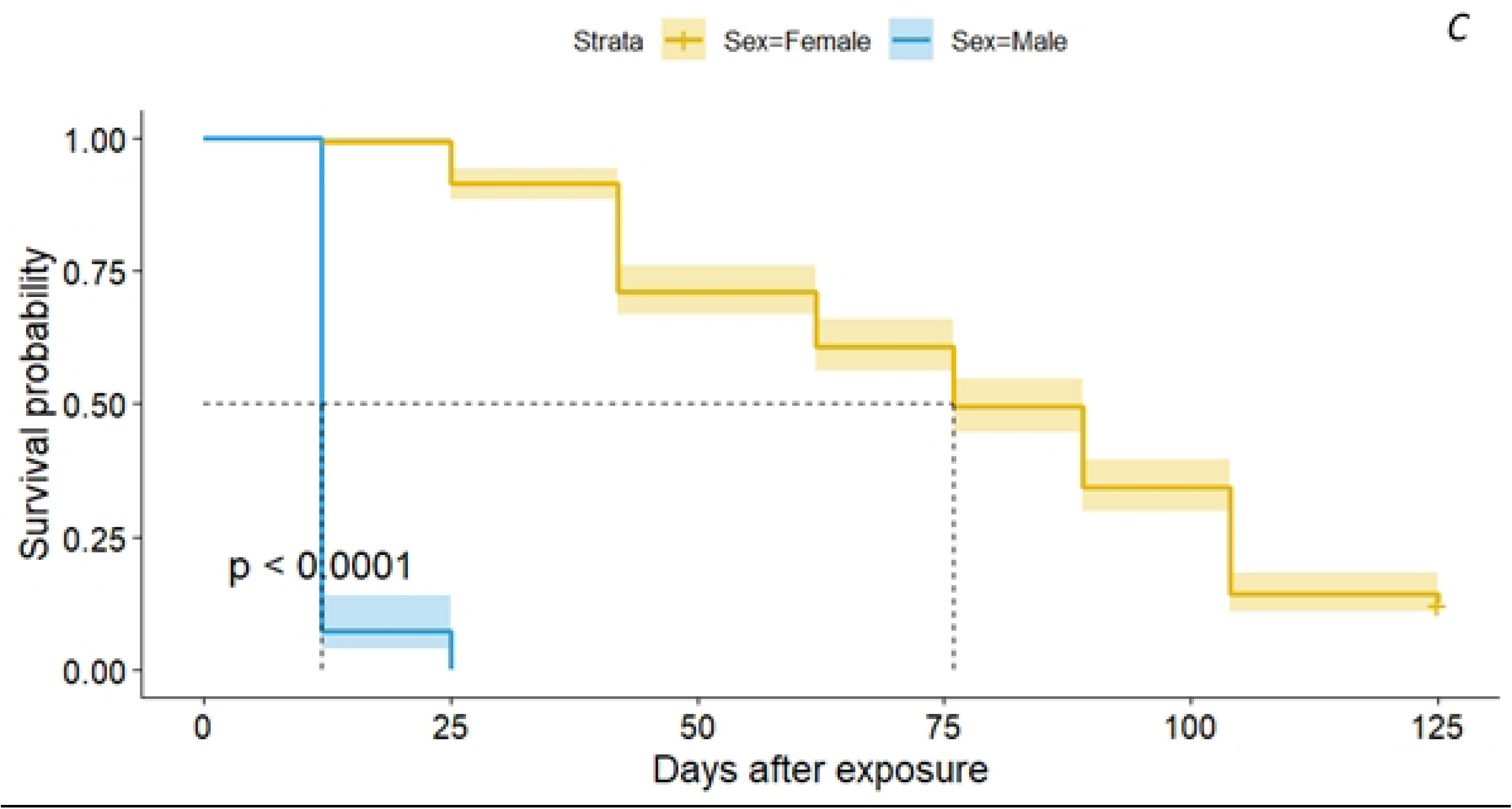
Kaplan Meier curves (with 95%CI) including log-rank test (shown p-value) for the survival of males and females in treatment A (Plain water+ organic water exposed to the wild on 1/10/2013), treatment **B** (Plain water only, exposed to the wild on 1/10/2013), and treatment C (Plain water only exposed to the wild on 15/10/2013). Black dashed lines represent the median survival for males {12 days) and females (76 days).

**Fig 8.**
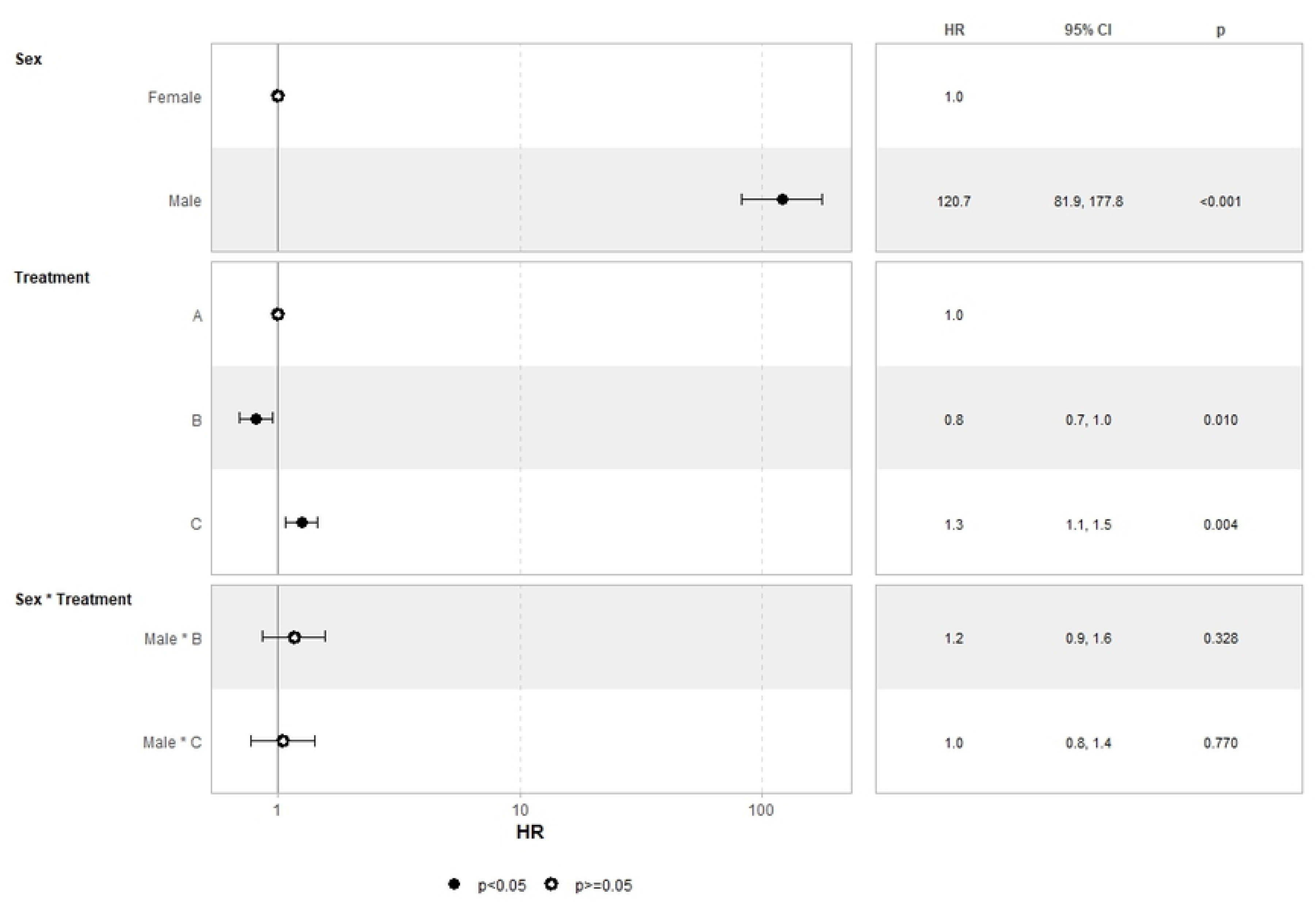
Forest plot depicting the hazard ratios (HR) for different sex, treatment, and interaction between sex and treatment groups exposed to Volos, during the winter 2012-13. Treatment A (Plain water, exposed to the wild on 1/10/2013), treatment **B** (Plain water + organic water, exposed to the wild on 1/10/2013), and treatment C (Plain water exposed to the wild on 15/10/2013).

## Discussion

The overwintering experiments of *Cx. pipiens f. pipiens* were carried out somewhat late in relation to the development of winter, but they are nevertheless informative of the survival of this species during the winter months in Central Greece. The successful females overwintering at all three regions, where our experiments were conducted, demonstrated that a significant proportion of the population was capable of surviving until next spring, constituting a remarkable basis for rapid growth of the species once temperatures allow. However, of particular interest is the fact that a proportion of the survived females managed to oviposit. To the contrary the males failed to survive until the end of the cold season, but they did live longer in Kalamaki than in Nea Anchialos. Given that males do not accumulate fat reserves, these differences are probably due to reduced metabolism due to the lower temperatures in Kalamaki, the mainland village, compared to Nea Anchialos, the coastal area. The presence of male Culex mosquitoes in early spring, given that male mosquitoes do not hibernate, is an indicator of when the first generation of mosquitoes, produced by post-diapause female Culex, achieves reproductive maturity [5].

The accurate prediction of WNV seasonal transmission cycles and the evaluation of the effectiveness of mosquito surveillance and control can be achieved by improving our understanding and knowledge of the initiation and termination of *Culex* diapause [5]. The two biotypes can hybridize, and hybrids show intermediate behaviour. Due to their more opportunistic feeding behaviour, hybrids are considered important bridge vectors which can transmit WNV from birds to humans [40–42]. In Europe the behavioural differences between the biotypes of *Cx. pipiens* may have an impact on their contribution to the WNV transmission cycle, therefore, it is essential to distinguish between biotypes when investigating the role of *Cx. pipiens* in WNV transmission [13,35]. In Germany, winter survival of WNV in vectors has been confirmed, suggesting its long-term persistence, as it has been detected several times in mosquitoes of the *Cx. pipiens* complex during the transmission period, and also in hibernating females of this complex in winter [38].

It is of vital importance for the survival of mosquitoes to select suitable hibernation sites, in which the microclimate can clearly have an impact on the winter survival of *Cx. pipiens f. pipiens*. The temperatures that are considered optimal for overwintering of adults are between 2 and 6°C [6]. Temperatures lower than 0°C can cause death after several days, whereas higher temperatures increase metabolic rates, which can result in the depletion of lipid reserves in female *Cx. pipiens f. pipiens.* For a better understanding of the dynamics of mosquito populations after winter and the way in which arboviruses survive in temperate regions, it would be of interest to explore which are the preferred hibernation sites and the determinants of winter survival in these sites [9,43]. Temperature plays an important role in whether mosquito vectors can overwinter in a given area, thus facilitating their establishment in new areas [44], whereas temperature variation can also affect disease transmission [45]. On the other hand, regarding the longevity of populations, Ciota et al. observed that the longevity of field populations tended to be longer than that of laboratory populations [46]. Under simulated field conditions, Abouzied noticed that female *Cx. pipiens* mosquitoes, in the winter/spring season survived for an average of 120 days, while in the summer/autumn season they survived for an average of 80 days, significantly exceeding the relative constant temperatures [47]. Data from the study of Spanoudis et al. indicated that certain biological parameters of *Cx. pipiens f. molestus* differ when measured at constant and fluctuating temperatures, highlighting the importance of testing fluctuating temperatures that simulate field conditions [48].

Understanding the environmental determinants of *Cx. pipiens* diversity is important because it can help us predict changes under future climate and land use regimes [49]. The longevity of adult *Cx. pipiens* has a negative correlation with temperature within the upper and lower survival limits. Warmer winters may expand the latitudinal zone within which *“molestus”* can survive aboveground, increasing hybridization between ecotypes in northern areas where they currently remain distinct. Such changes would be invisible at the morphological level, but nevertheless have potential consequences for disease transmission [37]. Climate change is considered to be a significant key factor that contributes to the global spread of mosquito-borne diseases. The rise of the median winter temperatures has been shown to extend host seeking and oviposition behavior in *Cx. pipiens* favouring overwintering of the virus in contrast to previous decades. Lower annual average winter and higher spring and summer temperatures have been linked to an increased risk of epidemic outbreaks [50,51]. Mosquito populations employ a diversity of overwintering mechanisms to better to withstand the consequences of climate change, such as sudden periods of unusually cold weather and decreased winter rainfall [51].

In the future, WNV and other flaviviruses that have significantly extended their distribution are predicted to become an increasing burden for public health systems [50]. The overwintering strategies could be used to design control approaches during the winter, as they would prevent the occurrence of high mosquito population in summer. A new approach to effectively control mosquitoes before they are capable of transmitting deadly pathogens to birds, humans and other animals, could potentially be provided by targeted pesticide applications in early spring [21,43]. The hibernating adults’ sites are of high importance and should be seriously examined. Microclimates encountered in urban areas (subways, houses) have often higher and more stable temperatures than outdoor environments [10]. Τhe experiment of Beleri et al. on the winter survival of adult *Ae. albopictus* in human made shelters, in Athens, Greece, revealed the importance of elaborating more on gaining further insights into winter survival of female mosquitoes [52]. Controlling the mosquito vector is an important control approach that should be adapted to the local vector ecology, considering the climatic conditions of microhabitats of overwintering vectors in urban areas [53–55].

The results of the current study could provide the basis for further research on the overwintering of mosquitoes in our country, covering a wider range of areas with different climatic conditions. Winter survival studies of native and invasive mosquito of medical significance will have specific importance for mitigating the disease and nuisance burdens caused by these mosquitoes. In northern temperate climates mosquitoes’ ability to successfully overwinter is to a large extent due to their ability to diapause [36]. A main part of these investigations should focus on areas where disease outbreaks occur repeatedly, with special emphasis on the detection of the WNV in overwintering mosquitoes of the genus *Culex* in order to elucidate the effect of this parameter on the maintenance of the infection each year and to mitigate the spread of the virus and reduce the impact of the upcoming epidemic.

## Supporting information

**S1 Fig.** Winter survival of *Culex pipiens f. pipiens* in Kalamaki (a mainland village) (A) and in Nea Anchialos (a coastal area) (B), Thessaly, 2012-2013.

**S2 Fig.** Kaplan Meier curves (with 95%CI) including log-rank test (shown p-value) for the survival of females in Nea Anchialos and Kalamaki. Black dashed lines represent the median survival for females in Nea Anchialos (66 days) and females in Kalamaki (67 days).

**S3 Fig.** Survival rate of *Culex pipiens f. pipiens* female adults regarding treatment 1 (a), treatment 2 (b), and treatment 3 (c), in each cage, in Volos, Thessaly, 2013-2014.

**S4 Fig.** Survival rate of *Culex pipiens f. pipiens*male adults regarding treatment 1 (a), treatment 2 (b), and treatment 3 (c), in each cage, in Volos, Thessaly, 2013-2014.

**S5 Fig.** Survival rate of *Culex pipiens f. pipiens* adults (a) females, (b) males, regarding the three treatments in Volos, Thessaly, 2013-2014.

## Author Contributions

C.I., C.H and N.T.P. conceived the study; N.T.P. and C.I. designed the experiments; C.I and P.T. collected the data, S.B., C.I. and E.V. analyzed the data; S.B., A.M., E.P., and N.T.P. wrote the first draft of the manuscript; N.T.P, C.H., C.I., S.B., E.P. and A.M. edited the manuscript. All authors reviewed the manuscript. C.H., E.P. A.M. and N.T.P acquired funding.

